# Attenuation of Hyperalgesia and Allodynia by some Phenolic Acids in Paclitaxel Induced Neuropathy

**DOI:** 10.1101/2021.01.17.427045

**Authors:** Shubhangi H. Pawar, Aman B. Upaganlawar, Chandrashekhar D. Upasani

**Affiliations:** Department of Pharmacology, MGVs Pharmacy College, Nashik, Maharashtra, India; Department of Pharmacology, SNJBs SSDJ College of Pharmacy, Chandwad, Maharashtra, India

**Keywords:** Hyperalgesia, Allodynia, Phenolic acids, neuropathy

## Abstract

Paclitaxel, an anticancer drug induced neuropathy is widely used animal model to evaluate new drugs in neuropathy. As oxidative stress is key contributor in pathogenesis of neuropathy, many phenolic acids with antioxidant potential are proven as neuroprotective. So,present work undertaken to evaluate effect of syringic acid and sinapic acid in paclitaxel induced neuropathy. We evaluated effect on mechanical and thermal hyperalgesia and allodynia which are vital signs of neuropathy. 4 weeks treatment by different doses of syringic acid and sinapic acid shown significant protective effect on hyperalgesia and allodynia in dose dependant manner assessed by Randello Selitto, hot plate, cold plate and Von Frey filament test. As these phenolic acids attenuates hyperalgesia and allodynia in neuropathy, can be therapeutically used in combination with current treatment of neuropathy.

**Summary statement:** Hyperlagesia and allodynia are major signs of neuropathy and this article focus on reduction of hyperalgesia and allodynia by syringic and sinapic acid in neuropathy induced by paclitaxel.

## Introduction

Cancer is a major contributor of death rate worldwide. Currently, modern chemotherapy is emerging to treat various complicated malignancies, which is increasing survival rate in cancer. But along with this, these chemotherapeutic agents are resulting into many adverse effects (Shahrak et.al., 2020). Taxanes are first-line group of antineoplastic drugs used to treat breast, lung, prostate and other gynecological malignancies. Paclitaxel is a prototypic semisynthetic drug from taxane group, derived from the precursor 10-deacetylbaccatin III. This precursor is obtained from tree *Taxus baccata*. Peripheral neurotoxicity and myelosuppression are two major toxicities associated with use of paclitaxel (Velasco and Bruna, 2015).

Mechanism of neuropathy induced by paclitaxel is thought to be dysfunctioning of microtubules in dorsal root ganglia, axons and Schwann cells. Paclitaxel forms abnormal microtubule bundles in the cytoplasm, which disrupts normal cell physiology and leads to cell proliferation. This impairs normal neuronal development and results in neuronal death (Scripture et.al., 2006). Paclitaxel, if given intraperitoneally, can produce a dose-limiting painful peripheral neuropathy and can be used as an animal model to evaluate the effect of various drugs in neuropathic pain (Saha et.al., 2012). Low doses of paclitaxel lead to dysfunction of axons and Schwann cells and produce pain hypersensitivity including allodynia and hyperalgesia, characterized by numbness, paresthesias and a burning pain in the hands and feet (Aswar and Patil, 2016). Peripheral nerve lesions may generate spontaneous pain and exaggerated responses to light touch i.e. allodynia and to temperature stimuli i.e. thermal hyperalgesia (Chaplan et.al., 1994).

Neuropathic pain is commonly observed adverse effect of chemotherapeutic agents which may affect patient’s life quality. Recently, many natural products are getting research attention due to their protective biological and pharmacological actions (Shahrak et.al., 2020). Various plants and their phyto-constituents are selectively studied in the treatment of neuropathy in rats (Ozyurt et.al., 2006). Phenolic acids are polyphenols, having anti-inflammatory and free radical scavenging action, have been proven as neuroprotective (Szwajgier et.al., 2017). In accordance with these effects of various phenolic acids, unravelled members of this class can be evaluated through rational research plan. Syringic acid (SY) is useful in treatment of diabetes, cardiovascular diseases, cancer and cerebral ischemia. It is having anti-oxidant, anti-microbial, anti-inflammatory, neuro-protective and hepato-protective activities. It effectively scavenges free radicals and reduces oxidative stress markers (Cheemanapalli et.al., 2018). Sinapic acid (SP) is widely used in pharmaceutical and cosmetic industries because of its potent antioxidant, anti-inflammatory, preservative and antimicrobial activity (Raish et.al., 2019). SP is effectively proven to prevent memory loss, counterbalancing oxidative stress and beneficial in the treatment of Alzheimer’s disease (Shahmohamady et.al., 2018). In accordance with this literature survey, the present study was undertaken to evaluate effect of SY and SP on hyperalgesia and allodynia in neuropathy.

## Material and Methods

Syringic acid and sinapic acid purchased from sigma Aldrich, USA. Standard drug gabapentin was supplied by Sun Pharma, India.

Research proposal was prepared as per guidelines of CPCSEA and granted approval by Institutional Animal Ethical Committee (IAEC) of SNJB’s SSDJ College of Pharmacy, Chandwad, India (CPCSEA approval letter No. SSDJ/IAEC/2018/01).

From the literature survey, minimum therapeutic doses of Syringic acid (SY) were finalized as 12.5, 25, 50 mg/kg/day and of Sinapic acid (SP) as 5, 10, 20 mg/kg/day orally (Li et.al.,2018, Shahmohamady et.al., 2018). Standard drug gabapentin 300 mg/kg/day p.o. was used to compare the results.

Animals used were Wistar rats of either sex and divided into 9 groups (*n=6*) and treated for 4 weeks as followings

1. Negative Control: received vehicle only.
2. Positive Control: Paclitaxel (Pcl) 2 mg/kg i.p. on four alternate days.
3. SY1: Pcl + Syringic acid 12.5mg/kg/day
4. SY2: Pcl+ Syringic acid 25mg/kg/day
5. SY3: Pcl+ Syringic acid 50 mg/kg/day
6. SP1: Pcl+ Sinapic acid 5mg/kg/day
7. SP2: Pcl+ Sinapic acid 10mg/kg/day
8. SP3: Pcl + Sinapic acid 20 mg/kg/day
9. Std: Pcl+Gabapentin 300 mg/kg/day p.o.

### Induction of neuropathy by paclitaxel

Clinically formulated paclitaxel solution for infusion was diluted with 0.9% sterile saline to achieve a 2 mg/ml solution for injection. Animals were dosed with 2 mg/kg paclitaxel or equivalent volume of vehicle solution intraperitoneally on four alternate days (0, 2, 4, and 6). Animals were dosed according to their weight (1 ml/kg) and were immediately returned to their home cages (Grifiths et.al., 2018).

Test drug treatment started from 6^th^ day of this injection schedule, considered as 0 week and treatment continued for next 4 weeks.

### Behavioural study

Peripheral nerve injury in neuropathy is characterized by behavioural biomarkers such as dysesthesia, hyperalgesia, allodynia and with motor inco-ordination (Wang and Wang, 2003)

#### a) Mechanical Hyperalgesia (Randall sellitto method)

Randall Selitto test is commonly used for testing acute mechanical sensitivity, measured by paw withdrawal threshold. Through the dome-shaped plastic tip of this apparatus, steadily increasing pressure applied on dorsal surface of the rat’s hind paw. The withdrawal threshold (in % CBK) for each paw recorded. Measurements repeated 2 or 3 times on each paw (Hanore 2006). Animal gently hold to immobilize it and its paw placed on the apparatus and the tip of device allowed to apply on paw with application of increasing mechanical force and withdrawal latency to the pressure supported was noted down (Khan et.al., 2019).

#### b) Heat hyperalgesia (Hot plate test)

Eddy’s hot-plate used to study the thermal nociceptive threshold by keeping the temperature at 55± 2°C. Animal individually tested by placing on the hot plate and paw licking latency (sec.) recorded. Test cut-off time of 20 sec was maintained (Krishnamurthy et.al., 2018).

#### c) Cold Allodynia (Cold plate test)

Here, the animal placed on the cooled plate at 5°C and the time to induce nociceptive behaviour indicated by shivering and paw licking recorded as the response time (Furgula et.al., 2018).

#### d) Mechanical Allodynia (Von Frey test)

Individually, rat placed on elevated maze in acrylic cage and adopted for test environment for at least 15min. From below the mesh floor, Von Frey filament applied to the planter aspect of hind paw. Enough force of filament applied against paw (causing slight bending) and hold for sec. Withdrawal of paw considered as a positive response (Sanklecha et.al., 2017) Observations of mechanical allodynia noted with 6 repeated application of varying force of Von Frey filament. Observations recorded in the format of OXXOXO, where O indicated- no withdrawal response and X indicates withdrawal response. This used method of observation is up and down method of Dixon and 50% gm threshold calculated by formula

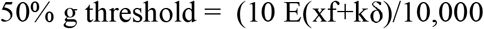

where Xf is log units of the last von Frey filament used; k is a tabular value for the pattern of positive/negative responses; and δ is a mean difference (in log units) between stimuli (here, 0.224) (Chaplan et.al., 1994).

## Results

### 1. Mechanical hyperalgesia (Randall Selitto test)

Withdrawal threshold of paw pressure is measured in terms of % CBK by Randall-Selitto apparatus. % CBK is found to be decreased significantly in positive control group indicating decreased paw withdrawal threshold and production of hyperlagesia compared to normal group. 4 weeks treatment with SY and SP has shown statistically significant (*p*<0.01**) protective effect indicted by significant increase in %CBK. Protective effect on hyperalgesia is dose dependant (SY 1, SY 2, SP 1, SP 2 protected with *p*<0.01* and SY 3, SP 3, Std with *p*<0.01**). All observations compared with positive control and standard gabapentin (Fig.1).

**Figure 1:**
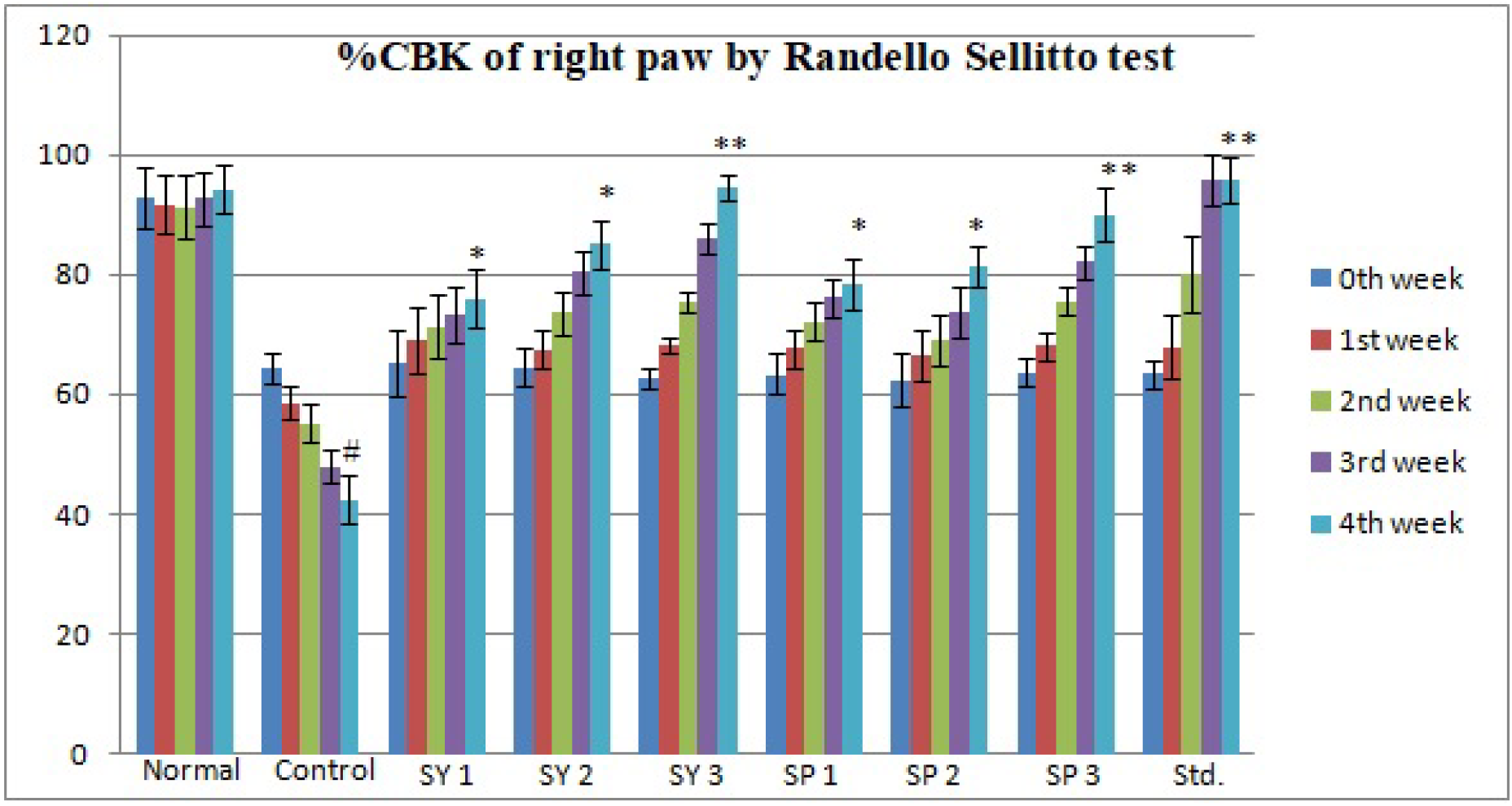
% CBK of right paw by Randello Sellitto test.

### 2. Heat hyperalgesia (Hot plate test)

Paw withdrawal latency and jumping response observed after placing animal on preheated plate (55°C). Thermal hyperalgesia is found to be produced in positive control group from 1^st^ week of paclitaxel injection. Heat produced hyperalgesia was significant on 4^th^ week of paclitaxel injection, indicating significant decrease in paw withdrawal latency compared to normal animals.Heat hyperalgesia is found to be prevented significantly after 4 weeks treatment with SY 1, SP 1 and SP 2 with *p*<0.01** and SY 2, SY 3. SP 3 and Std has shown statistically significant protection with *p*<0.01** (Fig 2).

**Figure 2:**
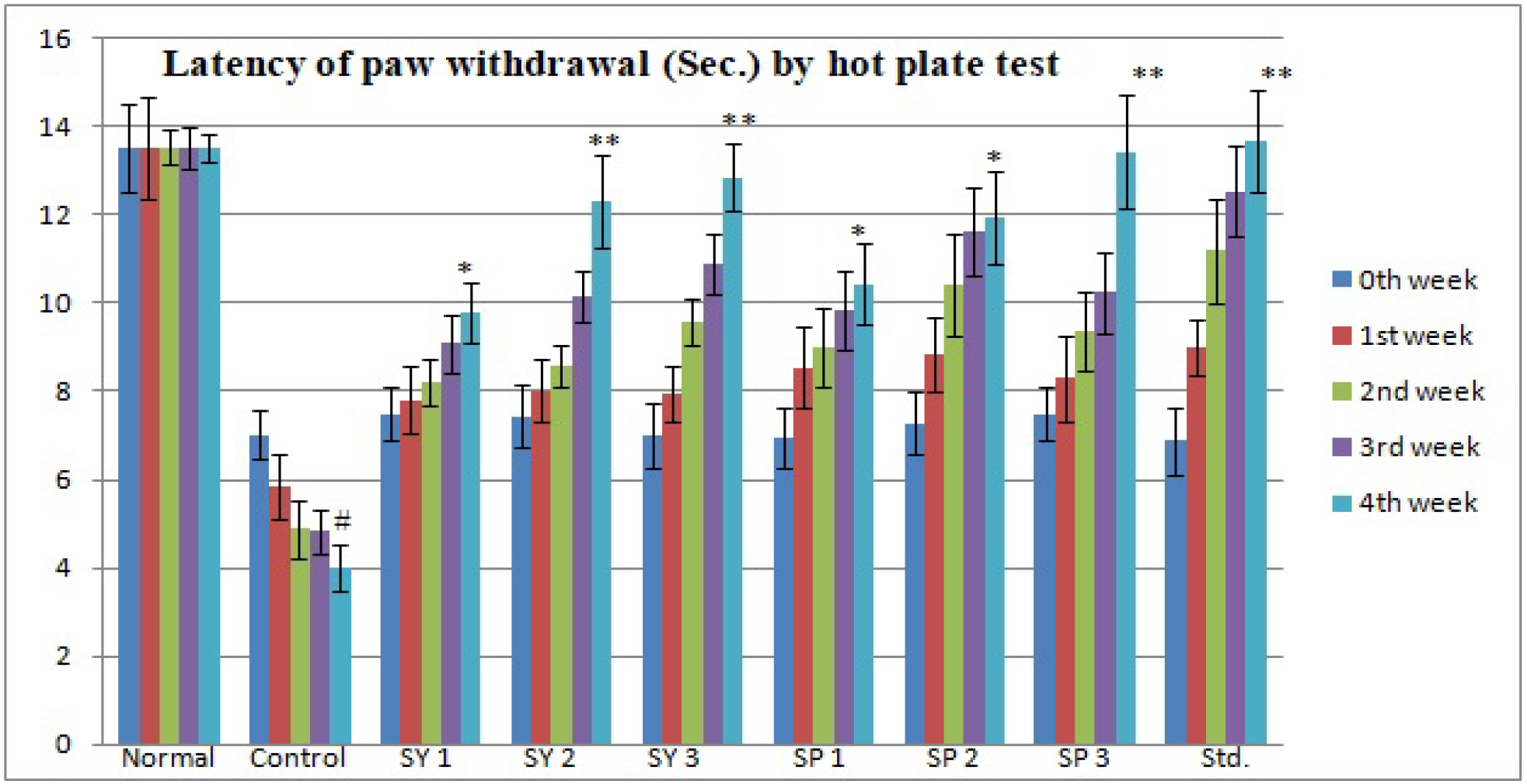
Latency (in sec.) of paw withdrawal by hot plate test.

### 3. Cold Allodynia (Cold plate test)

Time of onset of shivering from cold plate (5°C) is found to be decreased significantly in control group than normal group. Symptoms of cold allodynia observed from 1^st^ week of paclitaxel injection. In SY and SP decreased cold allodynia in dose dependent manner. These test drugs have shown protective effect from 2^nd^ week of treatment and 4 weeks treatment by SY 1, SP 1, SP 2 shown statistically significant protection of cold allodynia with *p*<0.05*. SY 2, SY 3,SP 3 and std drug (Gabapentin) shown it with *p*<0.01** (Fig.3).

**Figure 3:**
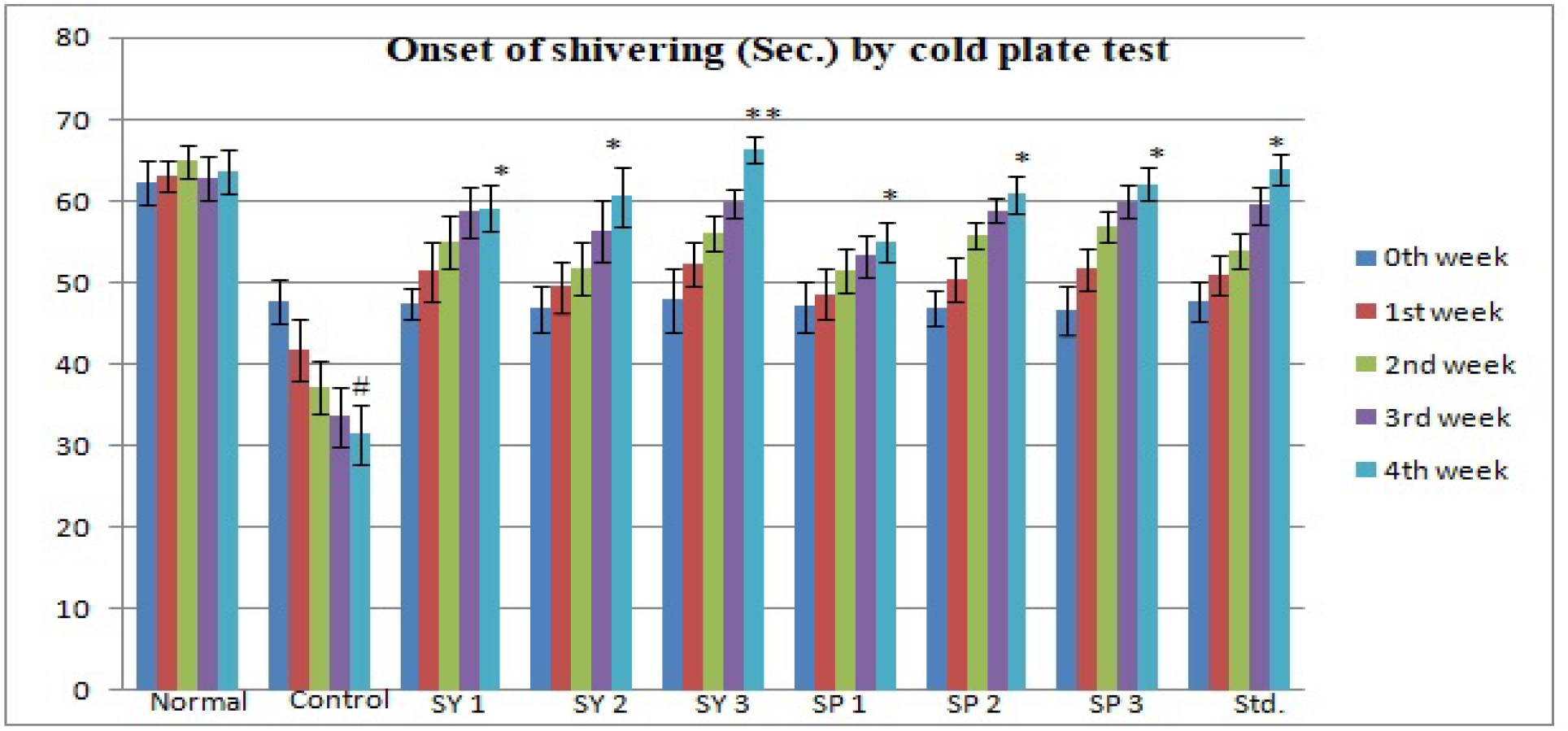
Onset of shivering (Sec.) by cold plate test.

### 4. Mechanical allodynia (Von Frey test)

Allodynia term refers to pain due to normally non noxious stimuli. It is observed in control group that allodynia produced from 1^st^ week of paclitaxel injection indicated by withdrawal response to minimum force of filament and thus significant decrease in 50% gm threshold. Treatment with all doses of SY and SP and Std have shown to increase force giving withdrawal response and 50% gm threshold in dose dependent manner (*p<0.01*\*\*) by changing observation format of Dixon up and down method (Fig.4).

**Figure 4:**
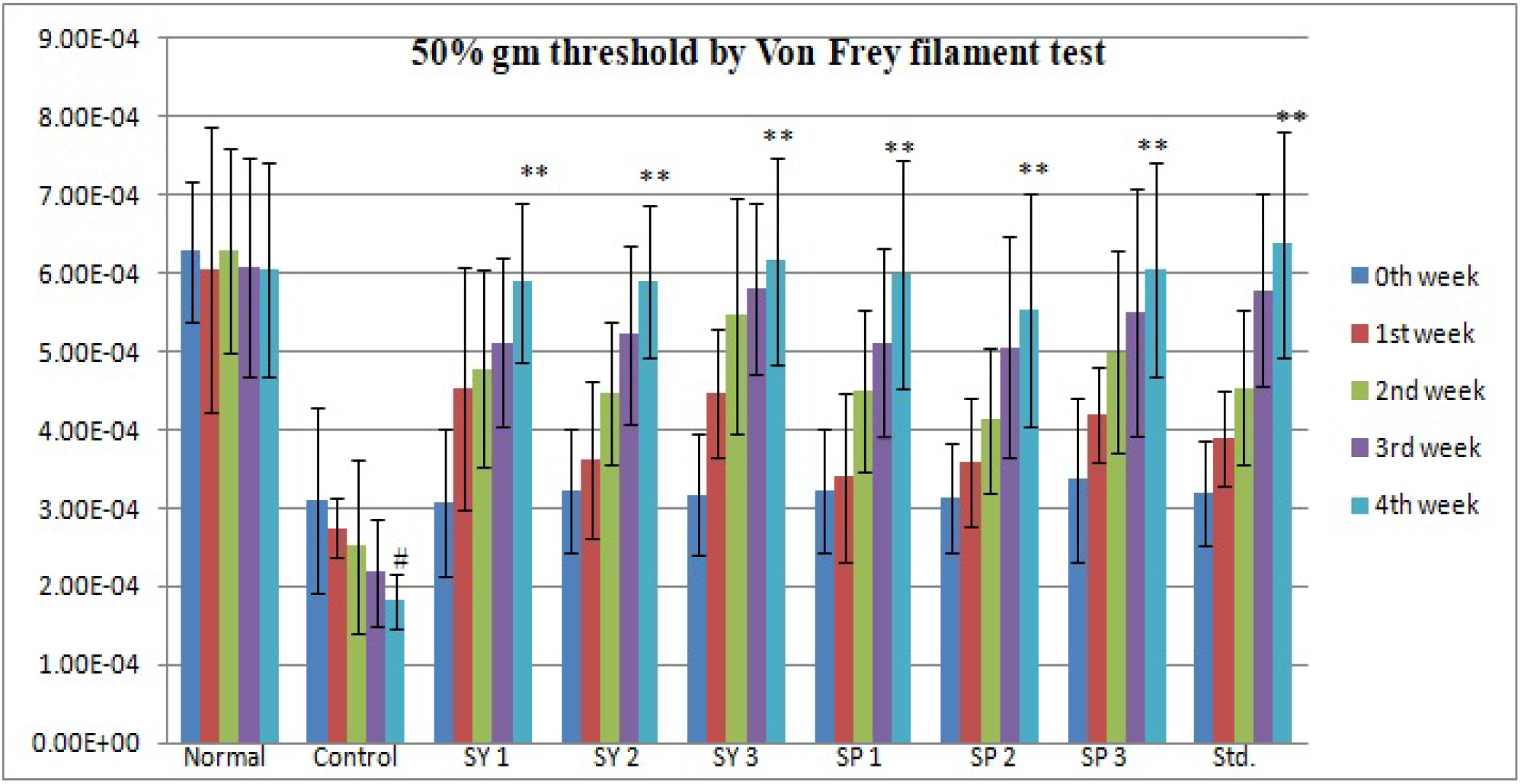
50% gm threshold by Von Frey filament test.

## Discussion

Paclitaxel, an anticancer drug induces neuropathy in dose dependant manner through dysfunctioning of microtubules in dorsal root ganglia, axons and Schwann cells and thus mechanism is useful to use it as effective animal model of neuropathy (Scripture et.al., 2006, Saha et.al., 2012). Neuropathic pain is mainly characterized by sensory abnormalities such as abnormal unpleasant sensation i.e.dysesthesia, increased intensity of response to painful stimuli i.e. hyperalgesia and pain in response to normally painless stimuli i.e. allodynia (Sausa et.al., 2016). At lower doses, paclitaxel lead to dysfunction of axons and Schwann cells and produce pain hypersensitivity including allodynia and hyperalgesia, characterized by numbness, paresthesias and a burning pain in the hands and feet (Aswar and Patil, 2016). In the present study, mechanical hyperalgesia is assessed by Randall Selitto test. The withdrawal threshold after mechanical pressure was considered to be associated with oxidative stress.^[31]^ Significant increase in withdrawal threshold indicated by increase in % CBK, which may be associated with reduction in oxidative stress by SY and SP. Syringic acid and Sinapic acid are proven potent antioxidants (Cheemanapalli et.al., 2018, Raish et.al., 2019). There are many evidences convincing role of antioxidants in treatment of neuropathic pain (Aswar and Patil, 2016). Thermo-nociception is evaluated by measuring withdrawal latency against fixed temperature. Heating surface allows stimulation of A or C thermonociceptors in rats, resulting in decreased pain threshold and withdrawal latency (Yalcin et.al., 2009). SY and SP have shown to increase time of paw withdrawal in hot plate test in dose dependant manner which may be associated with decreased stimulation of A or C fibers. The shivering and paw withdrawal response observed on the cold plate may be mediated by various factors. Cold stimuli activate neuronal Na^+^ channels and causes release of neurotransmitters mediated through pre-synaptic Ca^++^ channels. These ion channels are specifically localized on small, un-myelinated C-fibres and activation of these channels release neuropeptides, such as substance P and calcitonin gene related peptide from C-fibres. Thus, allodynia is produced through C-fibres and skin cooling activates thermoreceptors expressed in Ad fibres (Sachi et.al., 2008). This cold allodynia was found to be attenuated with SY and SP treatment in dose dependant manner.

In animal models of neuropathy, nociceptive behaviour can be provoked by minimum force of Von Frey filaments to the paw (maximum up-to 15 g). 50% threshold for nociceptive withdrawal latency provides quantitative assessment of mechanical allodynia expressed by von Frey filaments. SY and SP have shown significant increase in 50 % gm threshold in Von Frey filament test. Nitrosative stress, polymerase activation, increased excitability of ganglion neurons are factors thought to be associated with allodynia (Gao and Zheng, 2014).Thus in present study neuropathy symptoms are observed after 1 week of paclitaxel injection and 4 weeks treatment with SY and SP has shown protective effect.

### Conclusion

Syringic acid (SY) and Sinapic acid (SP) have shown attenuation in allodynia and hyperalgesia assessed by Randello Selitto, hot plate, cold plate and Von Frey filament test. This effect may be associated with their antioxidant, anti-hyperalgesic and anti-inflammatory effect. So these natural phenolic acids syringic acid and Sinapic acid can be therapeutically used in combination with current treatment of neuropathy.

## Acknowledgements

Authors are grateful to management trust of MGV, Nashik and SNJB, Chandwad for providing necessary laboratory provisions.

